# Electron transport chain RNAi in glutamate neurons extends life span, increases sleep, and decreases locomotor activity in *Drosophila melanogaster*

**DOI:** 10.1101/2023.01.08.523160

**Authors:** Jessie E. Landis, Kevin Sungu, Hannah Sipe, Jeffrey M. Copeland

**Author notes:** Corresponding author, (JMC).

## Abstract

RNAi targeting the electron transport chain has been proven to prolong life span in many different species, and experiments specifically with *Drosophila melanogaster* and *Caenorhabditis elegans* have shown a distinct role for neurons. To determine which subset of neurons is implicated in this life span extension, we used the GAL4/UAS system to activate RNAi against genes of Complex I and Complex V. We found life span extension of 18 – 24% with two glutamate neuron (*D42* and *VGlut*) GAL4 lines. We used the GAL80 system to determine if the overlapping set of glutamate neurons in these two GAL4 lines imparts the life span extension. Limiting GAL4 activity to non-*VGlut* glutamate neurons in the *D42* background failed to extend life span, suggesting that glutamate neurons have a unique role in aging. Interestingly, RNAi of the electron transport chain in *D42* glutamate neurons also caused an increase in daytime and nighttime sleep and a decrease in nighttime locomotor activity. Changes to sleep patterns and prolonged life span were not accompanied by any changes in female fertility or response to starvation. Our findings demonstrate that a small subset of neurons can control life span, and further studies exploring the role of the electron transport chain in aging can be focused on the activity of glutamate neurons.

## Introduction

Aging is marked by progressive tissue dysfunction, with common hallmarks in mammals, flies, worms, and fish. Such tissue dysfunction includes neurological impairments in cognitive function and memory and an increased likelihood of neurodegenerative diseases (1). Within cells, aging manifests as mitochondrial dysfunction, telomere loss, cell senescence, stem cell exhaustion, and disrupted cell signaling (2).

Mitochondrial dysfunction is a critical determinant in physiological aging, and the electron transport chain (ETC) is essential to mitochondrial activity. The ETC is comprised of five large protein complexes and produces most of the ATP and reactive oxygen species (ROS) in the cell. Changes by RNAi or mutations in individual components of the ETC are known to extend life span in many disparate species (3). Loss of ETC activity is not the only means to prolong life span. Overexpression of Ndi-1, a single component acting as NADH dehydrogenase/ubiquinone oxidoreductase I, can likewise extend life span in *Drosophila* (4,5). One connection between aging, ETC function, and life span is the production of ROS in the mitochondria. It has been observed that low concentrations of ROS in the cell stimulates mitohormesis, an adaptive cytoprotective response, but high concentrations accelerate aging (6,7). The concentration of ROS in particular is determined by the individual ETC component involved; a highly reduced pool of ubiquinone drives reverse electron transport and can prolong life span (8). Even though targeting ETC components by RNAi is sufficient to extend life span, the production of mitochondrial ROS has yet to be measured in *Drosophila* and *C. elegans* with activated RNAi.

While the creation and the cellular response to ROS are critical for aging, so is the removal of dysfunctional mitochondria. With advancing age, the number of dysfunctional mitochondria accumulate within a cell, with the older mitochondria characterized by their swollen size, fragmented appearance, and decreased respiratory ability (9). Concurrent with the increase of faulty mitochondria is the increased response of mitophagy. Mitophagy is one type of autophagy in which dysfunctional mitochondria are selectively sequestered and degraded by the lysosome. Several studies have uncovered that elevated levels of mitochondrial fission extend life span through the recovery of dysfunctional mitochondria (10,11). All of these studies underscore the importance of healthy mitochondria in promoting functional cellular activity and normal health span.

Particular subsets of cells can have cell non-autonomous effects for the aging of the entire animal. For example, RNAi manipulations of the ETC only in neurons can extend life span in both *Drosophila* and *C. elegans* (12,13). Once again, the importance of mitochondria in neuronal aging is not determined by activity of the ETC alone as changes in the proteins involved in mitophagy show similar effect (10,11). In addition, changes to mitochondrial fission in neurons has cell non-autonomous effects elsewhere. For example, upregulation of the mitophagy protein BNIP3 reduces age-related protein aggregation in muscles and improves intestinal barrier function (11). Certainly not all neurons affect life span equally. Activated RNAi of the ETC in dopamine and serotonin neurons shortens life span in *Drosophila* (14). Given the cell non-autonomous effects that neurons have on aging, questions remain as to which particular neurons impart these changes.

Glutamate neurons are involved in several behaviors impacted by age, like movement, sleep, and circadian rhythms. A decline in motor activity corresponds with advanced aging, and a few studies in *C. elegans* and *Drosophila* have shown that prolonged life span can negatively affect movement (15,16). Likewise, sleep patterns change with increasing age in both humans and flies, as observed specifically in fragmented and weakened sleep:wake cycles (17,18). Though sleep is a complex behavior involving many neuron clusters in the adult brain, glutamate neurons, specifically the dorsal clock neurons (DN1s), lateral posterior neurons (LPNs), and superior lateral protocerebrum neurons (SPNs), have been shown to have sleep promoting effects (19–21). Given these connections between life span, movement, sleep, and glutamate neurons, we activated RNAi against two components of the ETC in glutamate neurons and explored the effects on life span and these behaviors. Our results demonstrate that RNAi modification of the ETC in glutamate neurons extends life span, decreases nighttime locomotor activity, and increases daytime and nighttime sleep in flies.

## Materials and methods

### Fly strains used

The *D42-GAL4* (BL-8816), *VGlut-GAL4* (OK371; BL-26160), *ChAT-GAL4* (BL-6798), and *VGlut-GAL80* (BL-58448) fly strains were obtained from the Bloomington Stock Center (NIH P40OD018537, Bloomington, IN). The RNAi lines targeting the electron transport chain genes *ND-20/CG9172* (23256) and *ATPsynβL/CG5389* (22112) were purchased from the Vienna *Drosophila* RNAi Center (Vienna, Austria) and were backcrossed ten times to the *white^1118^* laboratory strain to minimize hybrid vigor (22).

### Life span assays

Flies for the life span assays were kept in a temperature-controlled (25 °C) container with 12:12 light:dark cycle at an average density of 25 flies per vial. Male and female adult flies were maintained separately. Flies were maintained on standard cornmeal food and flipped to fresh food twice a week. Prism9 (GraphPad, San Diego, CA) and Excel (Microsoft) software were used to analyze the life span data.

### Activity monitor

10 day old males with *D42-GAL4* activated RNAi against *CG9172* and *CG5389* or crossed with *white^1118^* laboratory strain were used for behavioral analysis. Flies were monitored with a 2 μW/cm^2^ intensity green LED light for 7 days at 12:12 light:dark cycle, followed by 7 days of 24 hrs dark cycle in Trikinetics activity monitors (Waltham, MA). Flies were maintained at 22°C, 60% relative humidity. 32 male flies were used per condition, and activity monitor tubes contained a plug of fly food at one end. Data were analyzed using the ShinyR-DAM program (23) and Microsoft Excel.

### Fertility

To measure female fertility in the *D42*-GAL4 activated RNAi lines, 3-day old virgin females were mated to 2-day old *w^1118^* males. Each fly cross contained 5 adult females and 7 adult males and each experiment had three vials. Parental flies were transferred to new vials 2-3 times per week, and progeny were counted every day after eclosion. Experiments were conducted in duplicate.

### Stress Resistance

Response to starvation were recorded in the *D42*-GAL4 activated electron transport chain RNAi lines. Starvation conditions were created by placing 10-day old females on 1% agar. Experiments were conducted in duplicate.

## Results

In previous studies, RNAi targeting individual components of the electron transport chain only to neurons was sufficient to prolong *Drosophila* life span by approximately 15%, a difference of approximately 10 days (12). Given these results, we set out to uncover which neuronal subtypes are important for life span extension. In this study, we used the GAL4/UAS system to activate RNAi in specific subsets of neurons. Specifically, the *D42*-GAL4 and the *VGlut*-GAL4 (*OK371*) driver lines were used to broadly target an overlapping set of glutamate neurons, and *ChAT-* GAL4 to target cholinergic neurons. Using both *D42-* and *VGlut-GAL4* driver lines was important as these lines show overlapping expression in adult and larval brains and in motor neurons. RNAi was used to affect the activity of the Complex I gene *CG9172* and the Complex V gene *CG5389*. The life span of flies with activated RNAi was compared to that of heterozygous GAL4 flies without the UAS-RNAi elements.

We observed a life span extension of 23% and 18 % (p < 0.0001), respectively, when RNAi targeted *CG9172* and *CG5389* in *D42*-GAL4 glutamate neurons (Fig 1A, B; Table 1). Similar life span extension of 24% and 22% (p < 0.0001) was observed using the *VGlut-GAL4* line (Fig 1C, D). A smaller life span extension of about 12% was observed with RNAi activation in cholinergic neurons (p < 0.0001). Maximum life span, as measured as the life span of the top 10% longest living cohort, was also prolonged in flies with RNAi activated by both glutamate GAL4 lines. A marginal increase in maximum life span was seen with RNAi in cholinergic neurons (Table 1). Although the response was not as consistent as for female flies, average male life span was prolonged by 11 to 28% in ETC RNAi activated by the *D42-* and *VGlut*-GAL4 lines, showing that the response is conserved among the sexes (S1 Fig; S1 Table). Life spans of unactivated UAS-*CG5389*-RNAi and UAS-*CG9172*-RNAi showed only a 5% increase when compared to a *white^1118^* control line, indicating that extension is not simply an artifact of the UAS-RNAi element itself (S2 Fig).

**Fig 1.**
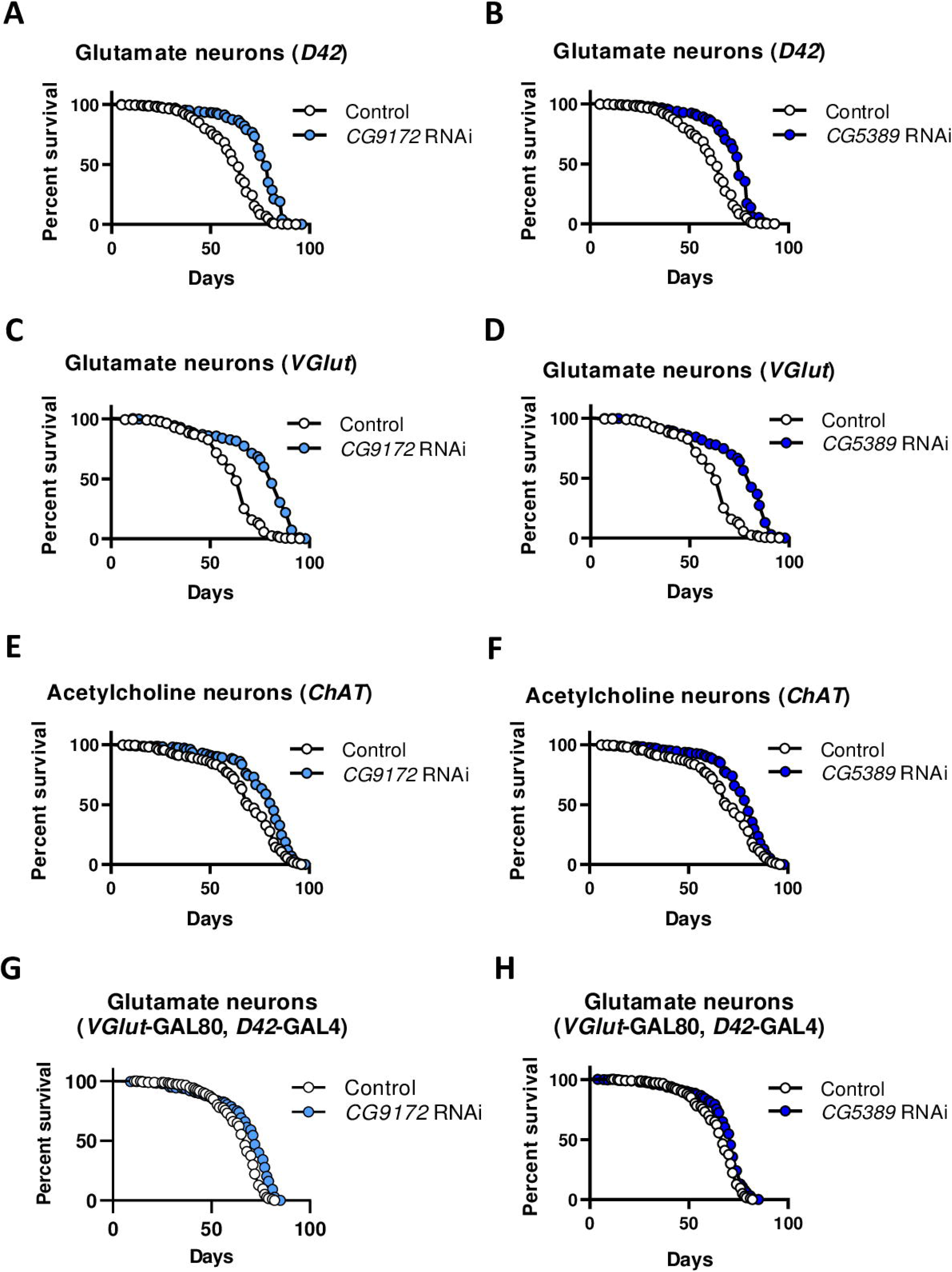
RNAi of the electron transport chain in glutamate neurons is sufficient for female life span extension in *Drosophila*. Females of the indicated GAL4 driver were mated to a *white^1118^* control (open circles) or UAS-*CG9172-*RNAi (light blue), or UAS-*CG5389*-RNAi (dark blue) lines. (A, B) Survival curves of RNAi against *CG9172* and *CG5389* RNAi in *D42*-GAL4 glutamate neurons show comparable 23% and 18% (p < 0.0001, log rank test) life span extension. (C, D) In *VGlut-GAL4* glutamate neurons, RNAi of *CG9172* and *CG5389* leads to an extension of 24% and 22%, respectively (p < 0.0001). (E, F) RNAi of *CG9172* and *CG5389* in *ChAT-GAL4* acetylcholine neurons extends life span by 12% (p < 0.0001). (G, H) Activating RNAi against *CG9172* and *CG5389* to the set of non-overlapping set of glutamate neurons (*D42*-GAL4; *VGlut*-GAL80) minimizes life span extension to 6% and 5%, respectively (p < 0.0001).

**Table 1.**
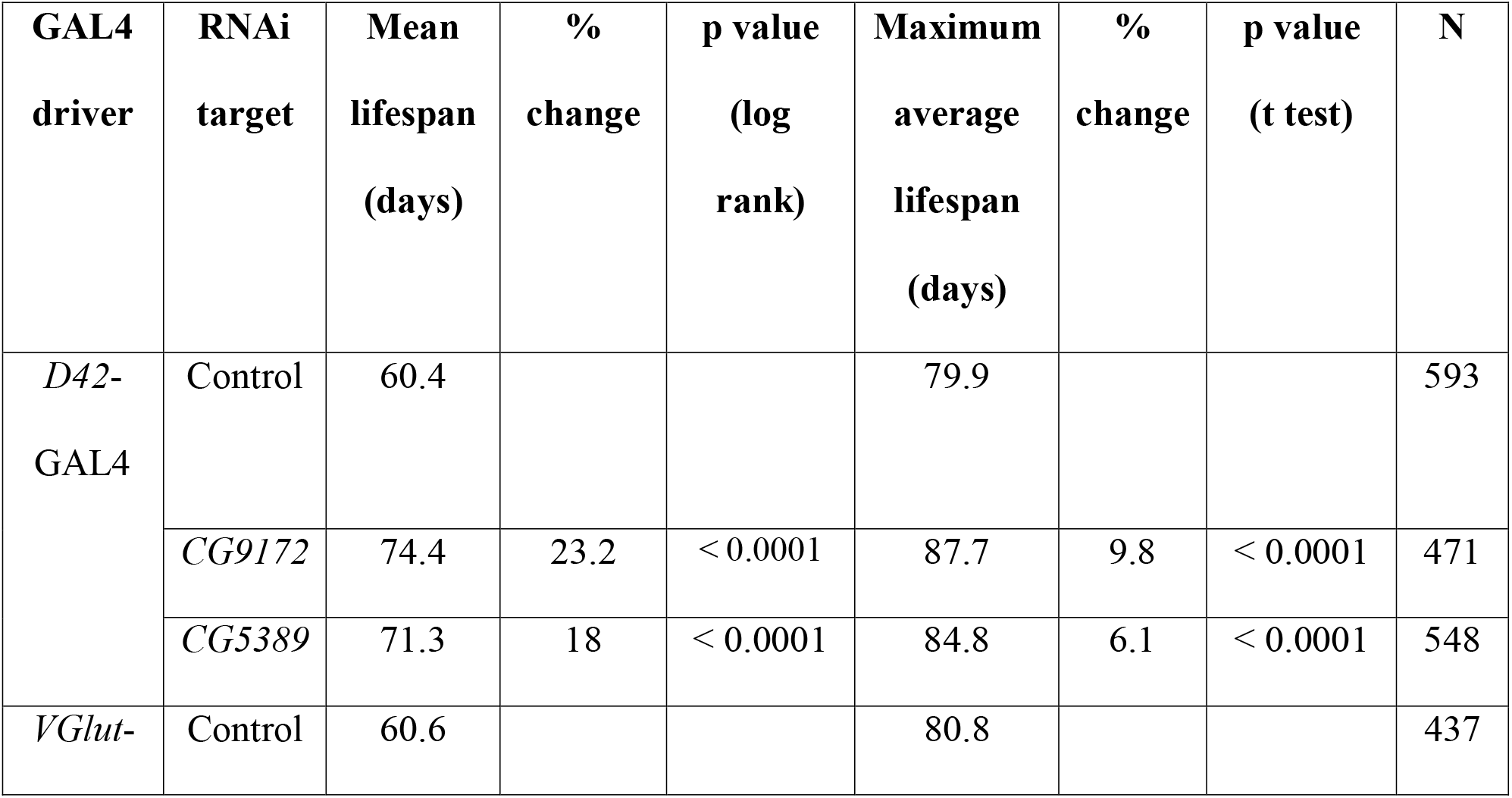
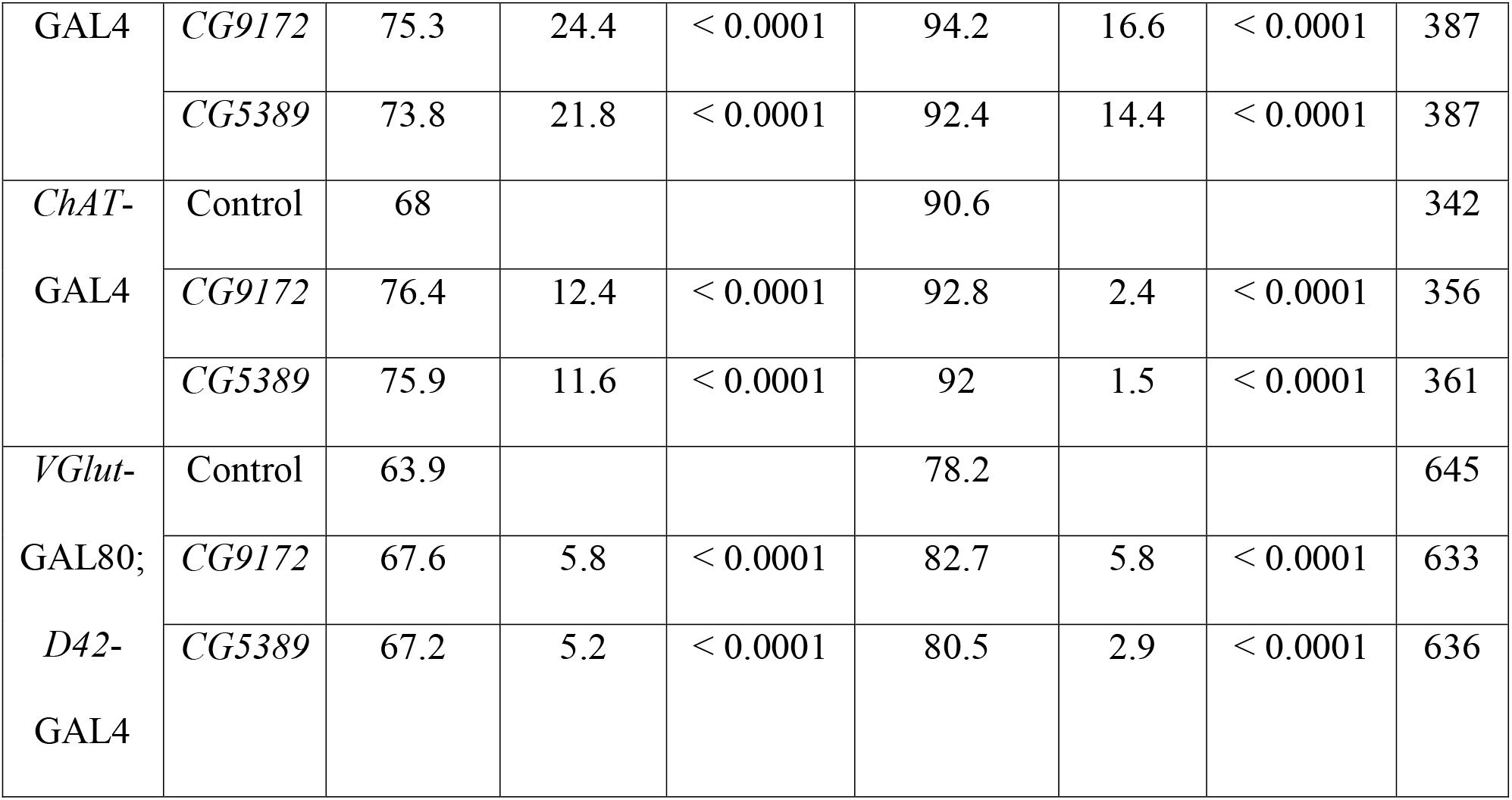
Descriptive statistics of female life span in neuron-specific RNAi of the electron transport chain.

To determine if the overlapping set of glutamate neurons are responsible for the life extension, the GAL4/GAL80 system was employed to block RNAi activation in these cells. In our experiment, ETC RNAi was activated in *D42*-GAL4 glutamate neurons, but inhibited in *VGlut*-GAL80 expressing neurons. In these flies, life span extension was significantly reduced in activated *CG9172* and *CG5389* RNAi lines to 5 – 6% (Fig 1G, H). One possible explanation for the remaining life span extension is residual GAL4 activity and RNAi activation despite the presence of the GAL80 inhibitor. Nevertheless, these results highlight the crucial role that glutamate neurons play in life span extension.

### Decreased *Drosophila* motor activity during dark phase in a 12 hour light/dark cycle

With such a noticeable life span extension from glutamate neurons with activated ETC RNAi, we wanted to explore a possible relationship between activity and aging. We used Trikinetics activity monitors (Waltham, MA) on flies with RNAi targeting *CG9172* and *CG5389*. Cohorts of 10 day old flies were placed in the activity monitor in 12:12 light/dark (LD) conditions for 7 days and then switched to all dark (DD) settings for 7 days. Overall average activity was measured for 24 hour periods, as well as activity specifically during the light phase and dark phase. As a control group, *D42*-GAL4 flies outcrossed to *w^1118^* were included.

*CG9172* and *CG5389* RNAi flies were significantly less active compared to the *D42*-GAL4 control flies. In the 24 hour LD cycle, *CG9172* RNAi flies were 32% less active and *CG5389* RNAi flies were 40% less active (p < 0.0001) (Fig 2; S2 Table). This decrease in activity was specific for nighttime. During the 12 hour dark phase, the *CG9172* RNAi and *CG5389* RNAi flies were 64% and 69% (p < 0.0001) less active than the control group. In the light phase, the decline in activity did not meet statistical significance.

**Fig 2.**
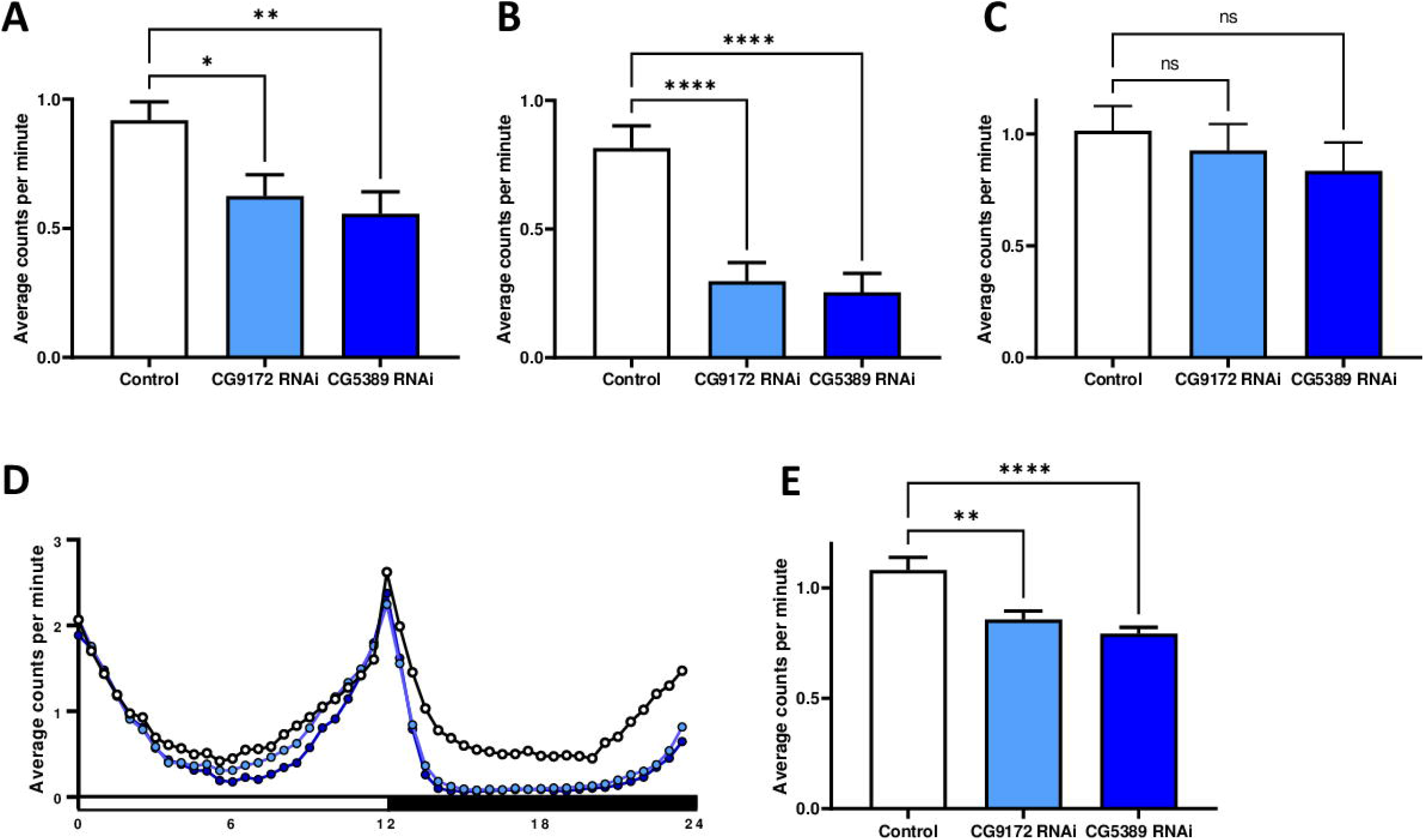
RNAi of the electron transport chain in glutamate neurons decreases nighttime activity in 10 day old flies. Adult male flies of either the *D42*-GAL4 control line (white) and activated *CG9172* RNAi (light blue) and *CG5389* RNAi (dark blue) were tested for their activity starting at 10 days of adulthood. Data are presented as means +/- SEM and compared using one-way ANOVA with a Tukey’s post-hoc test. (A) Average activity over 24 hours in a LD cycle saw a decrease of 32% (p = 0.029) and 39% (p = 0.005) in RNAi targeting *CG9172* and *CG5389*. (B) Targeting *CG9172* and *CG5389* by RNAi leads to a decrease in average activity during the dark phase of a 24 hour LD cycle by 64% and 69% (p < 0.0001), respectively. (C) Average activity during the light phase was not statistically altered in *CG9172* and *CG5389* RNAi activated flies (p = 0.86, p = 0.54), respectively. (D) Representative activity profile over 24 hours. (E) RNAi targeting of *CG9172* and *CG5389* in a 24 DD cycle decreased activity by 21% (p = 0.0011) and 32% (p < 0.0001), respectively.

In general, all flies tended to be more active in the DD setting than in the 24 hour LD phase, but the activated RNAi flies were consistently less active than the *D42*-GAL4 control line. In DD, the *CG9172* RNAi flies were 21% less active (p < 0.0011) and the *CG5389* RNAi flies were 32% less active (p < 0.0011).

### Increased sleep observed during a 12 hour light/dark cycle in electron transport chain RNAi flies

The most pronounced effect on activity in the ETC RNAi lines was during the dark phase in the LD conditions. Given the timing of this effect, we wanted to explore a possible connection with sleep. Sleep was defined as bouts of inactivity longer than 5 minutes (18). Using the data collected from the activity monitors, the average amount of sleep was calculated per fly for a 30 minute period. Data was also analyzed depending on the phase of the 24 hour light/dark cycle.

The data clearly shows a dramatic increase in the amount of sleep in the ETC RNAi flies. The average amount sleep for the 24 hour LD period increased by 95% and 115% (p < 0.0001) for the *CG9172* RNAi and *CG5389* RNAi lines, respectively (Fig 3; S3 Table). The increase in sleep was not limited to a particular phase of the LD cycle, but the changes during the dark phase (p < 0.0001) were more statistically significant than the light phase (p = 0.03 – 0.002). The activated RNAi flies also exhibited a significantly longer sleep bout length than the control. Bout length increased 31.1% and 41.8% (p < 0.0001) in the *CG9172* and *CG5389* RNAi lines.

**Fig 3.**
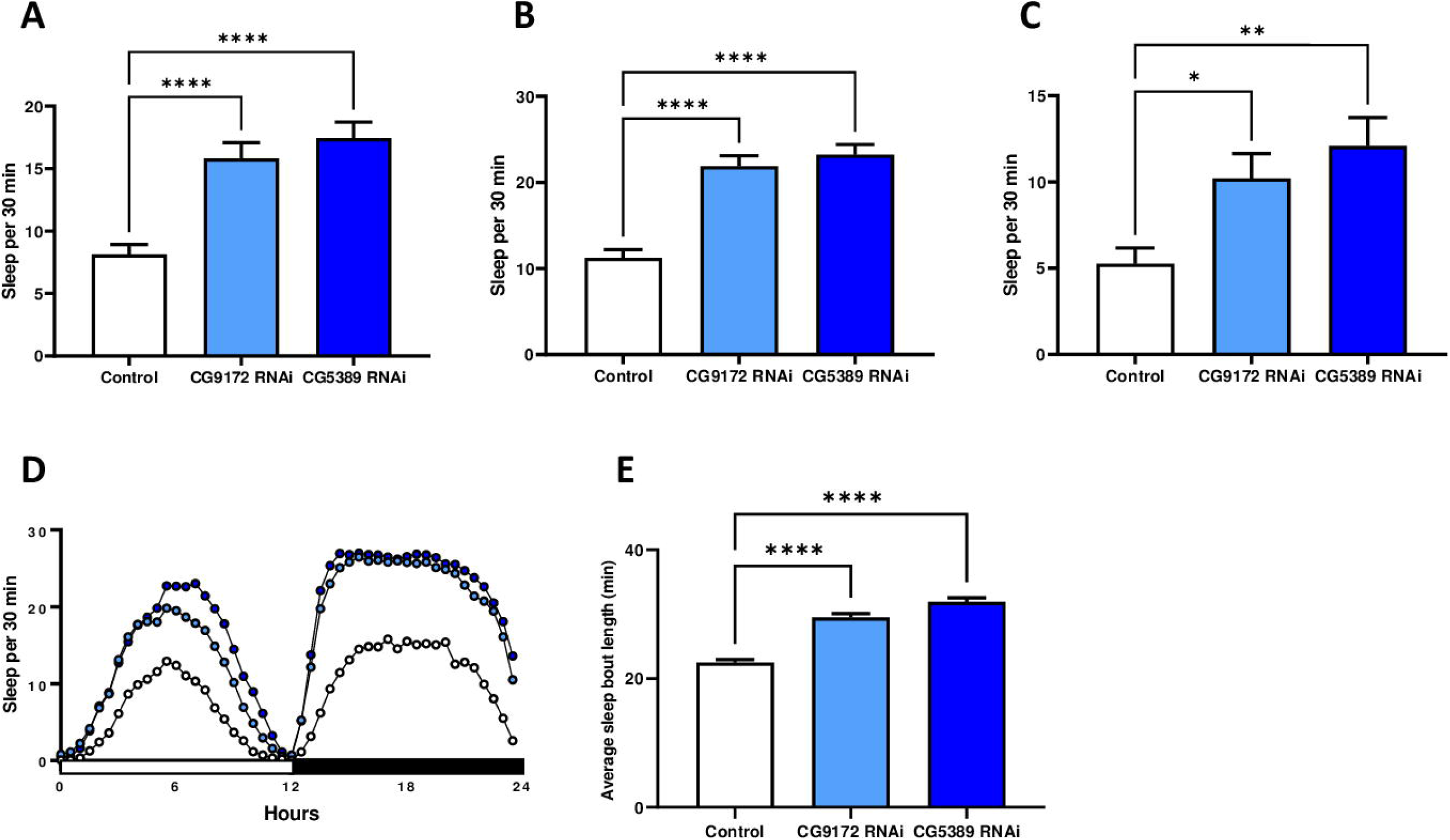
RNAi of the electron transport chain in glutamate neurons increases sleep in 10 day old flies. Adult male flies of either the *D42*-GAL4 control line (white) and activated *CG9172* RNAi (light blue) and *CG5389* RNAi (dark blue) were tested for overall sleep starting at 10 days of adulthood. Sleep data is presented as average time asleep per 30 minutes, shown as means +/- SEM, and compared using one-way ANOVA with a Tukey’s post-hoc test. (A) The average amount of sleep over 24 hours in a LD cycle increases by 95% (p < 0.0001) and 115% (p < 0.0001) in RNAi targeting *CG9172* and *CG5389*. (B) Average sleep amount during the nighttime phase increases by 95% and 107% (p < 0.0001). (C) A similar increase in sleep amounts is seen during the light phase for *CG9172* RNAi (94%, p = 0.035) and *CG5389* RNAi (130%, p = 0.0022) adults. (D) Representative sleep profile over 24 hours. (E) RNAi targeting of *CG9172* and *CG5389* increases length of a sleep bout by 31% (p < 0.0001) and 41% (p < 0.0001), respectively.

### No effect on female fertility and starvation with electron transport chain RNAi in glutamate neurons

In addition to monitoring activity and sleep, fertility rate and response to starvation conditions were studied to assess possible physiological impacts of ETC RNAi. Oftentimes, extended life span causes a reciprocal decline in reproductive capacity (24). For example, decrease in ETC functioning in *Caenorhabditis elegans* shows a decrease in fertility as well as movement (15,25). In this study, we measured female fertility by counting the amount of adult progeny produced over the first 30 days of adulthood in flies with RNAi activated in *D42*-GAL4 glutamate neurons. RNAi of *CG9172* or *CG5389* did not affect female fertility in comparison to the *D42*-GAL4 control line (Fig 4). These results are consistent with a previous study that noted no change in female fertility in pan-neuronal RNAi of the same electron transport chain genes (12). To extend the study of possible physiological costs for prolonged life span, we examined the ETC RNAi flies response to starvation. Starvation conditions were created by placing 10 day old females on 1% agar. ETC RNAi flies had life spans 4% shorter than the *D42*-GAL4 control lines, though these results were only statistically significant in the *CG5389*-RNAi flies. Given the minimal effect and the difference between the two ETC RNAi lines, we suggest that these results do not show any meaningful sensitization to starvation.

**Fig 4.**
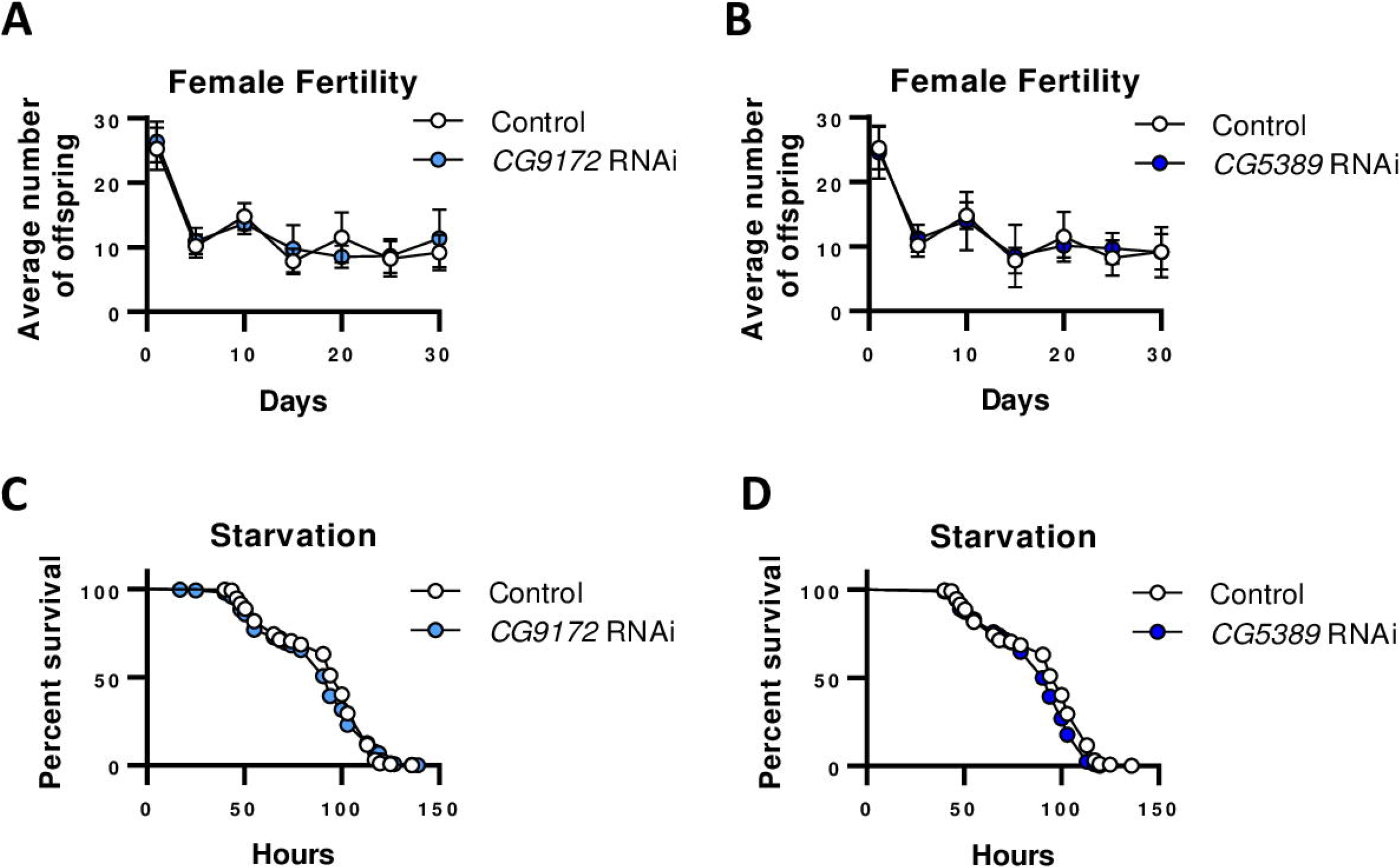
Activation of electron transport chain RNAi in glutamate neurons confers no significant effect on female fertility and starvation response. Females with the *D42*-GAL4 driver were mated to a *white*^1118^ control line (open circles) or RNAi targeting *CG9172* (light blue) or *CG5389* (dark blue) lines. (A, B) Activation of RNAi to *CG9172* (p = 0.98, two way ANOVA) and *CG5389* (p = 0.68) in glutamate neurons does not affect female fertility. (C, D). Starvation in females with activated RNAi to the electron transport chain genes specifically in *D42*-GAL4 neurons has a minimal effect on overall survival (*CG9172*, no change, p = 0.32; *CG5389*, decrease 4%, p = 0.0005).

## Discussion

In this report, we present evidence for a cell non-autonomous role for glutamate neurons in modulating *Drosophila* life span. RNAi targeting individual components of the ETC in glutamate neurons was sufficient to extend life span by 18 to 24%. RNAi of the same ETC components caused a remarkable increase in sleep and a decrease in activity specifically during the dark phase of a 24 LD cycle.

Previous work has established that RNAi of the ETC in neurons alone is sufficient to extend life span. Specifically, activation of RNAi in neurons of components from Complex I, IV, and V and ectopic expression of the single Complex I subunit Ndi1 prolongs life span (12,26). All neurons do not participate equally in aging, since ETC RNAi in dopamine and serotonin neurons shorten life span (14). The *D42*-GAL4 and *VGlut*-GAL4 are routinely used to drive gene expression in glutamate neurons and are known to be active in an overlapping set of cells (27,28). In our experiments, we notice a significantly smaller life span extension with the addition of the *VGlut-* GAL80 element to flies with RNAi active in *D42*-GAL4 glutamate neurons. These results strongly suggest that glutamate neurons are uniquely involved in life span extension.

The fact that alterations of mitochondrial function in neurons affect *Drosophila* life span in a cell non-autonomous manner is not surprising, as changes in the intestines have similar effects. Intestine-specific expression of *Ndi1* or RNAi inactivation of a Complex V subunit can significantly lengthen life span in flies and nematodes, respectively (5,25). Life span extension is not limited directly to the ETC as promoting mitochondrial biogenesis in the intestines had similar effects (29). It is worth noting the complex association between longevity and intestinal integrity. Dysfunction of the intestinal barrier precedes aging-related decline and can cause alterations of the microbiota, though changes to the microbiota are not driving life span (30,31). Our results might have connection with the demonstrated importance of the intestines in life span determination. Other studies have demonstrated that changes in the nutrient sensor AMPK in neurons can induce intestinal autophagy, decrease intestinal barrier dysfunction, and prolong life span (32). Glutamate motor neurons are known to innervate the digestive tract hindgut and proventriculus, potentially connecting these two aging-important tissues (33,34). It is of course important to determine the importance of intestinal integrity in life span extension observed in flies with glutamate neuron-specific ETC RNAi.

In this work, we identify the role of the electron transport chain in sleep and motor activity. We expected an overall decrease in fly activity with ETC RNAi targeting glutamate neurons, as excitatory motor neurons in flies are glutamatergic. It was surprising, however, that the effect on activity was limited to the dark phase in the LD cycle. Conversely, ETC RNAi drove the flies to sleep more throughout the 24 hr cycle, with longer sleep bouts to account for the increase in total sleep. Our results showing a genetic connection between sleep and aging join a significant body of research tying the two. In typical *Drosophila* aging, sleep becomes more disrupted with age, and younger flies have prolonged sleep bouts and more nighttime sleep (17). In our experiments, 10 day old flies with activated ETC RNAi slept much longer, similar to young flies carrying a Complex V point mutation (35). The effect of ETC RNAi on sleep during the aging process is still to be explored as our experiments only used 10 day old flies. Other studies have uncovered neuronal genes involved in both sleep and life span, with the genes affecting sleep and aging in a direct relationship (36). The fact that glutamate neurons play an important role in sleep is well established, but there are some conflicting studies about whether these neurons promote sleep or wakefulness (20,37,38).

Why does RNAi of the ETC in glutamate neurons affect sleep and longevity? It is well established, though counterintuitive, that genetic disruption of the ETC by RNAi does not affect overall ATP levels (12,25). Certainly, it is worth exploring what long term effect ETC RNAi has on mitochondrial activity in glutamate neurons and how the RNAi affects neuronal activity or age-related loss. One means to address possible changes to glutamate neuron activity is through the use of a heat-activated allele of the *dTrpA1* cation channel to over-activate glutamate neurons. Alternatively, the use of a temperature-sensitive *shibire* allele could inhibit glutamate neuron function and alter life span. Sangston and Hirsh have shown that the *TH*-GAL4 driver can change fly activity contingent on temperature sensitive *dTrpA1* expression (39).

It is important to note that the entire electron transport chain is involved in the aging process and sleep rather than just one subunit. RNAi targeting separate components of Complex I and Complex V had quite similar and robust effects. It is therefore unlikely that any one component would have an independent role outside of the ETC. Additionally, it is unlikely that the changes presented in this report are an artifact of RNAi off-targeting. The hairpin DNA sequences used to target *CG9172* and *CG5389* share no sequence similarity (Vienna Drosophila Resource Center, Vienna, Austria).

The role of the nervous system in regulating life span is certainly complex and each neuronal subtype does not contribute equally. The results presented here show that RNAi targeting the electron transport chain in glutamate neurons is sufficient to prolong life span. RNAi of the same electron transport chain components leads to a sharp increase in daytime and nighttime sleep and a specific decrease in nighttime activity. It is possible that aging and sleep are independent biological phenomena, but this report provides another genetic link between these processes. Given that only a small subset of neurons can influence such biologically important behaviors, understanding the precise role of the electron transport chain is of significant importance.

## Acknowledgments

The authors would like to thank Eli Wenger and Derek Harnish for their assistance with the life span experiments and Jay Hirsh (University of Virginia) for the use of his Trikinetics activity monitors. Stocks obtained from the Bloomington Drosophila Stock Center (NIH P40OD018537) were used in this study. J.M.C. is supported by a Small Project Research Grant from the Virginia Academy of Science and the Department of Biology at Eastern Mennonite University.

## Supporting information

**S1 Fig. RNAi of the electron transport chain in glutamate neurons extends life span in male *Drosophila*.** Males of the indicated GAL4 driver were mated to a *white*^1118^ control (open circles) or UAS-*CG9172-*RNAi (light blue), or UAS-*CG5389*-RNAi (dark blue) lines. (A, B) Survival curves of RNAi against *CG9172* and *CG5389* RNAi in *D42*-GAL4 glutamate neurons show comparable 11% and 17% (p < 0.0001, log rank test) life span extension. (C, D) In *VGlut*-GAL4 glutamate neurons, RNAi of *CG9172* and *CG5389* leads to an extension of 28% and 26%, respectively (p < 0.0001). (E, F) RNAi of *CG9172* and *CG5389* in *ChAT*-GAL4 acetylcholine neurons extends life span by 2% and 5%, respectively (p = 0.81, p = 54). (G, H) Activating RNAi against *CG9172* and *CG5389* to the set of non-overlapping set of glutamate neurons (*D42*-GAL4; *VGlut-GAL80*) minimizes life span extension to 2%, respectively (p = 0.91, p = 0.44).

**S2 Fig. Controls for the UAS-RNAi lines slightly extend life span in female *Drosophila*.** Females of the indicated UAS-RNAi lines were mated to a *white*^1118^ control (blue circles) and the *white*^1118^ control line was used for comparison (open circles). (A, B) Survival curves of outcrossed UAS-*CG9172*-RNAi (light blue) and UAS-*CG5389*-RNAi (dark blue) compared to the *white*^1118^ control both show 5% (p = 0.0011, p = 0.0041, respectively, log rank test) life span extension.

**S1 Table.**
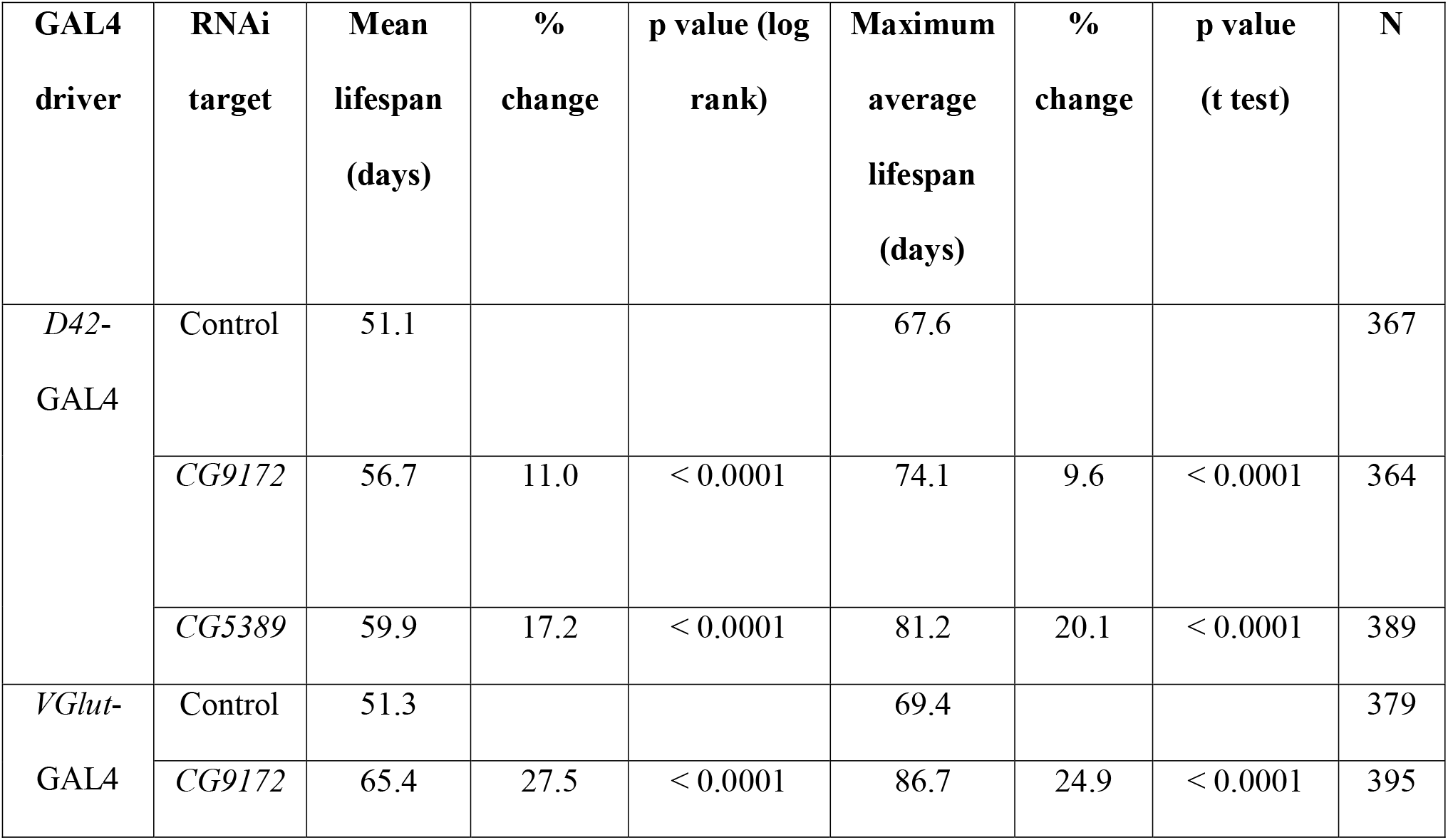
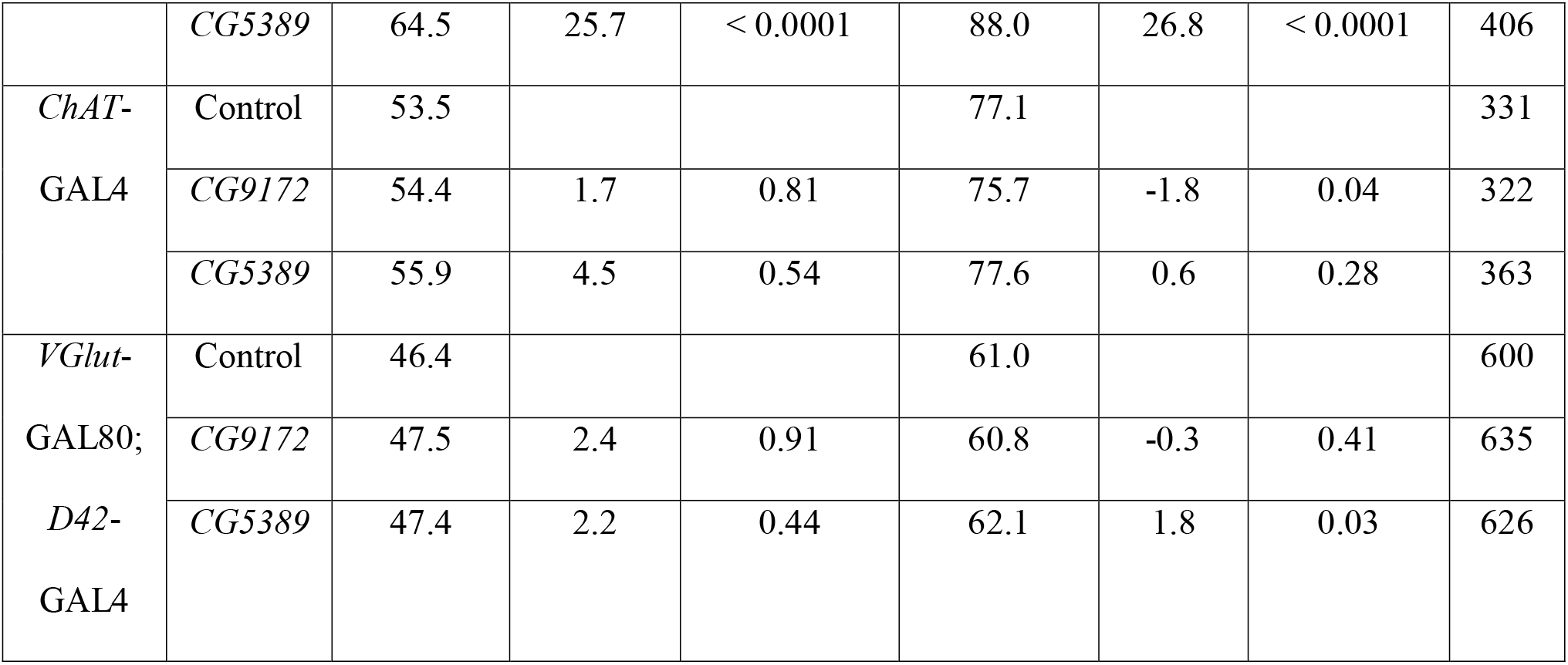
Descriptive statistics of male life span in neuron-specific RNAi of the electron transport chain.

**S2 Table.** Descriptive statistics of male motor activity in glutamate neuron (*D42*) specific RNAi of the electron transport chain.

|  | Genotype |  |  |
| --- | --- | --- | --- |
| Measurement | *D42-GAL4* / +<br>(control) | *D42>CG9172* RNAi | *D42>CG5389* RNAi |
| Total activity in LD<br>(avg count per min) | 0.9187 ± 0.072 | 0.62 ± 0.08 | 0.56 ± 0.09 |
| p values |  | 0.029 | 0.005 |
| % change |  | 32.10 | 39.50 |
| Activity at night, LD<br>(avg count per min) | 0.815 ± 0.09 | 0.296 ± 0.07 | 0.253 ± 0.07 |
| p values |  | <0.0001 | <0.0001 |
| % change |  | 63.681 | 68.957 |
| Activity during day,<br>LD (avg count per<br>min) | 1.014 ± 0.11 | 0.926 ± 0.12 | 0.835 ± 0.13 |
| p values |  | 0.86 | 0.539 |
| % change |  | 8.679 | 17.653 |
| Total activity in DD<br>(avg count per min) | 1.08 ± 0.06 | 0.857 ± 0.04 | 0.7931 ± 0.03 |
| p values |  | 0.0011 | <0.0001 |
| % change |  | 20.648 | 31.574 |

**S3 Table.**
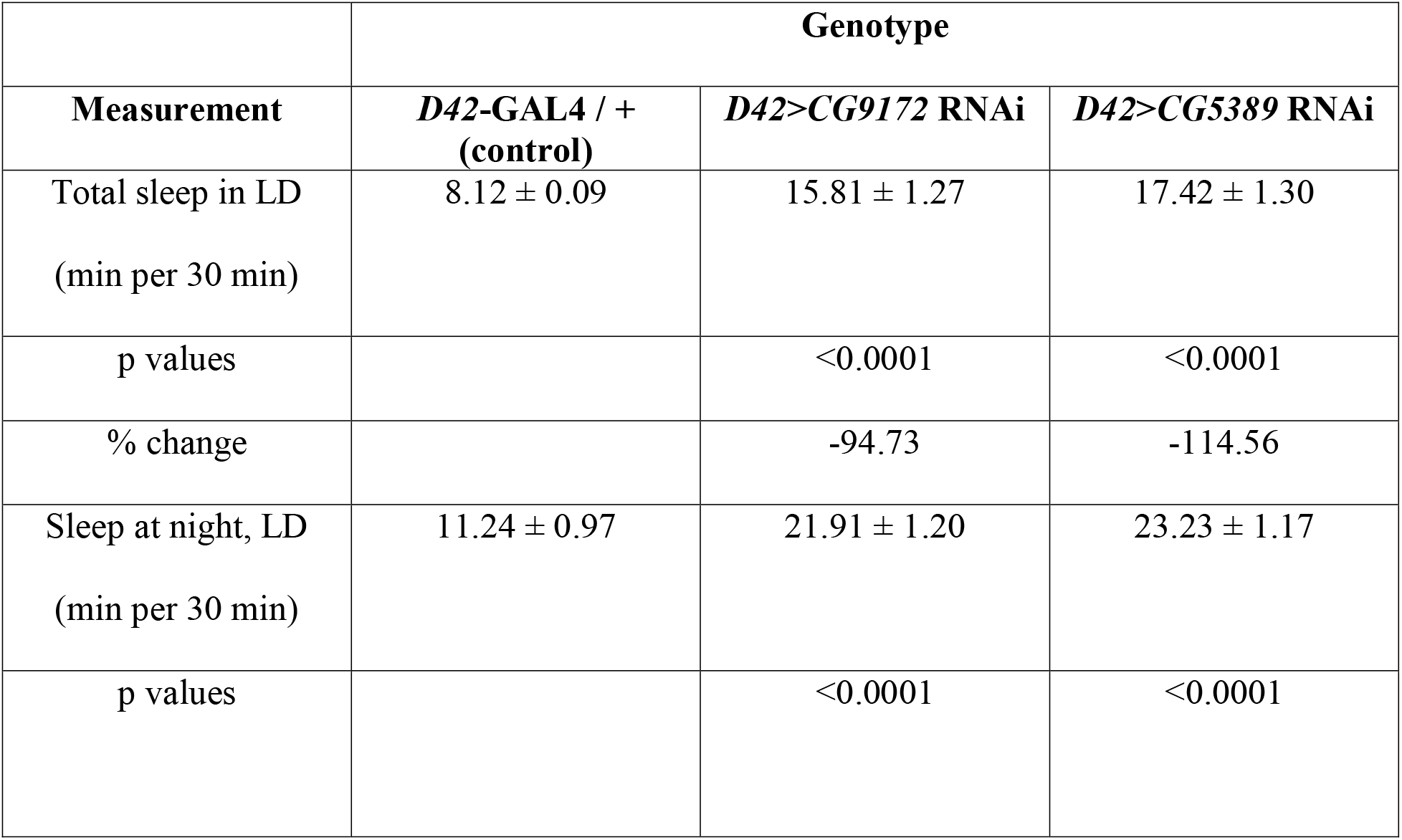
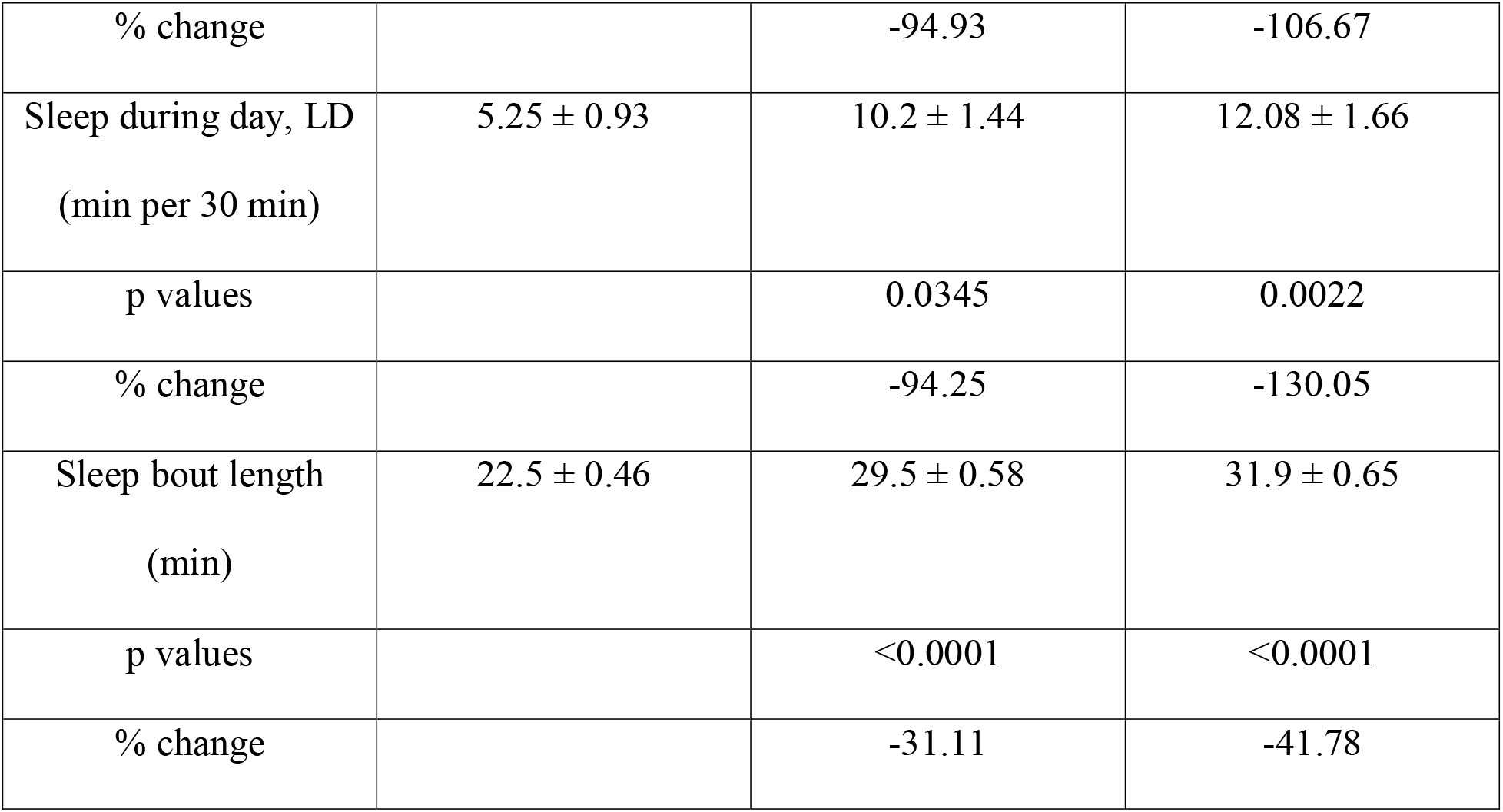
Descriptive statistics of male sleep in glutamate neuron (*D42*) specific RNAi of the electron transport chain.

## Notes

### Competing Interest Statement

The authors have declared no competing interest.

