## Supplementary figures and images for "Electron transport chain RNAi in glutamate neurons extends life span, increases sleep, and decreases locomotor activity in *Drosophila melanogaster*"

### S1 Fig

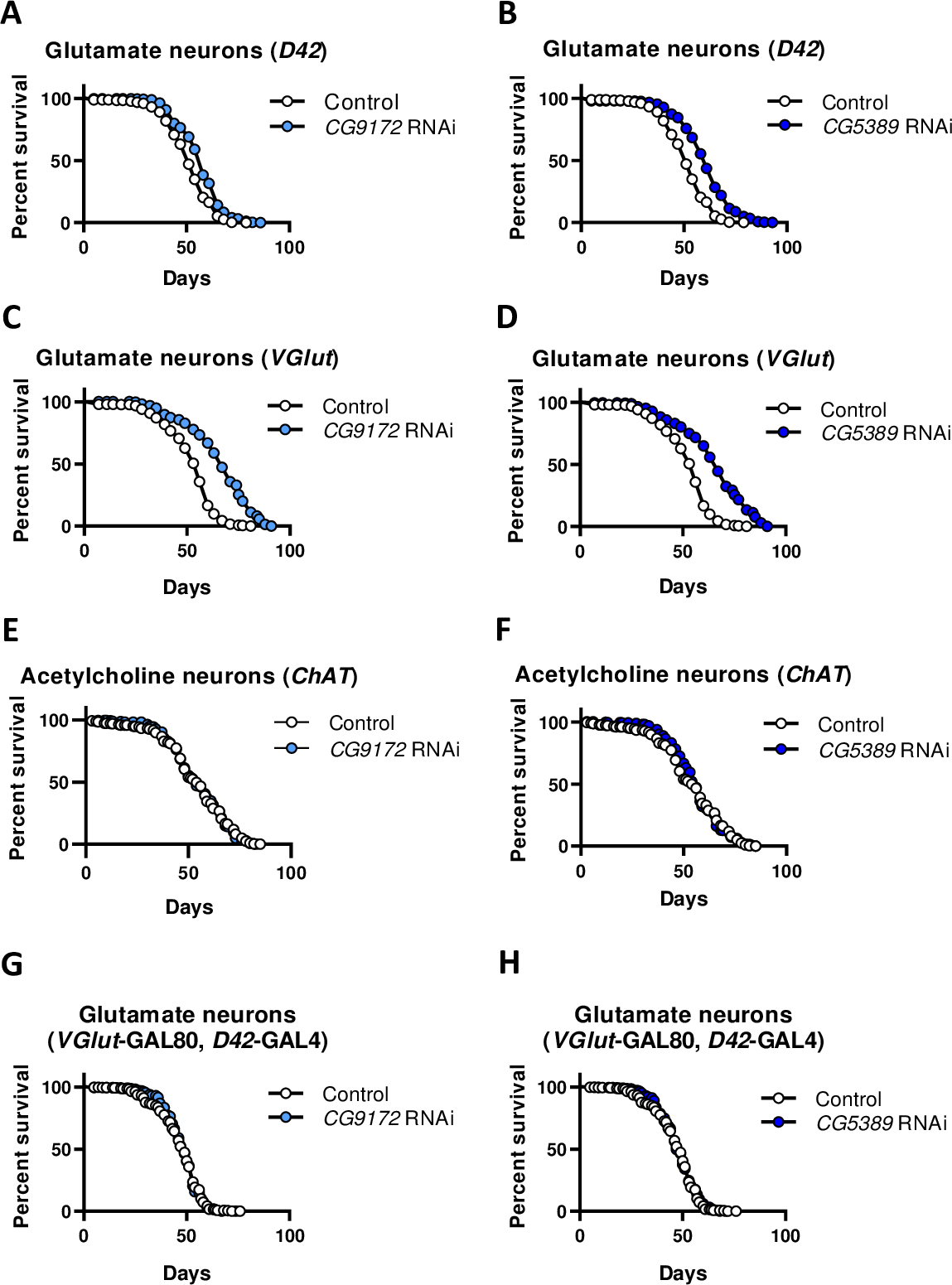

### S2 Fig

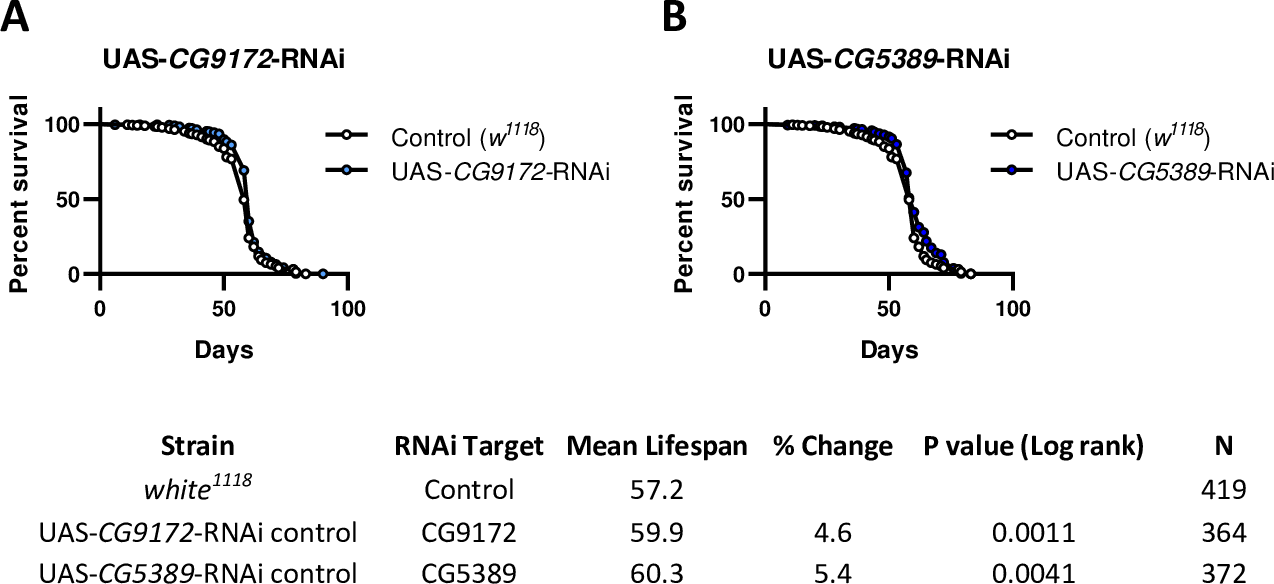
